# Influence of insertion sequences on population structure of phytopathogenic bacteria in the *Ralstonia solanacearum* species complex

**DOI:** 10.1101/2022.07.16.500299

**Authors:** Samuel TE Greenrod, Martina Stoycheva, John Elphinstone, Ville-Petri Friman

## Abstract

*Ralstonia solanacearum* species complex (RSSC) is a destructive group of plant pathogenic bacteria and the causative agent of bacterial wilt disease. Experimental studies have attributed RSSC virulence to insertion sequences (IS), transposable genetic elements which can both disrupt and activate host genes. Yet, the global diversity and distribution of RSSC IS are unknown. In this study, IS were bioinformatically identified in a diverse collection of 356 RSSC strains representing four phylogenetic lineages, and their diversity investigated based on genetic distance measures and comparisons with the ISFinder database. IS distributions were characterised using metadata on RSSC lineage classification and potential gene disruptions by IS were determined based on their proximity to coding sequences. In total, we found 24,732 IS belonging to eleven IS families and 26 IS subgroups, with over half of the IS found in the megaplasmid. While IS families were generally widespread across the RSSC phylogeny, IS subgroups showed strong lineage-specific distributions and genetically similar bacterial strains had similar IS contents. Further, IS present in multiple lineages were generally found in different genomic regions suggesting potential recent horizontal transfer. Finally, IS were found to disrupt many genes with predicted functions in virulence, stress tolerance, and metabolism, suggesting that they might be adaptive. This study highlights that RSSC insertion sequences track the evolution of their bacterial hosts, potentially contributing to both intra- and inter-lineage genetic diversity.

## Introduction

Genetic variation is the raw material for selection and evolutionary change. In bacteria, genomic variation is generated by replication errors resulting in single nucleotide polymorphisms (SNPs), small and large genome rearrangements, and gene gain and loss via horizontal gene transfer (1). Further modifications that affect gene expression include epigenetic modifications via DNA methylation and gene disruptions caused by the movement of mobile genetic elements including integrated bacteriophages (prophages) and insertion sequences (IS). IS are small transposable elements which can move within a single genome or horizontally between different cells, generating within and between population genetic variation. IS have two main structural components: a transposase, the enzyme responsible for IS translocation; and transposase-flanking terminal inverted repeats, linked to transposase binding, DNA cleavage, and strand transfer (2). IS are highly diverse and are classified into families based on the gene sequence of their transposase, and further into subgroups based on the presence and order of transposase protein domains (for review see (3)). In contrast to transposons, IS seldom carry auxiliary genes that would have effects on host bacterial fitness (3). Instead, the impact of IS on host fitness is dependent on their genomic location. Most fitness-associated IS insertions are found within coding sequences, resulting in deactivation of particular genes. IS-mediated gene disruptions are prevalent across many bacterial taxa and have been found to alter bacterial traits including antimicrobial resistance (4–6), virulence (7–9), and metabolism (10,11). Moreover, IS can also alter host gene expression by inserting close to genes, which can result in gene promoter disruptions or even increased neighbouring gene expression as some IS contain either entire or partial promoter regions. Examples leading to differential gene expression caused by IS insertions include increased host resistance to antibiotics (12,13) and phages (14), and activation of virulence (15) and metabolic pathways (16,17).

While IS have been studied in both eukaryotic host-associated and free-living bacteria, most of the research on IS host fitness effects has focused on human bacterial pathogens (for review see: (18)) and only a few studies have been published on plant pathogenic bacteria. In the rice pathogen *Xanthomonas oryzaei*, IS have been shown to inactivate genes in the *gum* cluster responsible for the biosynthesis of extracellular polysaccharides (19), in addition to disrupting the virulence-related *purH* gene (20). Moreover, IS virulence gene disruptions have been found in *Pseudomonas syringae* pv. *phaseolicola*, responsible for halo blight of the common bean, via IS insertion into a potential avirulence gene (21). Notably, in the same genera, IS have also been linked to the horizontal transfer of virulence genes (22–24), suggesting they likely play a role in both elevated and reduced host virulence. Recently, IS content was also investigated in the plant pathogenic bacterial *Ralstonia solanacearum* species complex (RSSC), a causative agent of bacterial wilt disease, through a genomic analysis of 62 complete RSSC genome sequences in the NCBI database (25). This study identified 20 IS families, including some which were widespread across the bacterial phylogeny and others which were only found in specific host phylogenetic lineages. IS were often located nearby, or inserted into, genes with potential roles in virulence, resistance to oxidative stress, and toxin production, potentially affecting the fitness of their hosts (25). These findings supported previous analyses of RSSC IS which identified disruptions in type III effectors (26) and in the global virulence regulator phcA (27), the latter of which resulted in spontaneous phenotypic conversion between non-virulent and virulent pathogen genotypes. It was also recently shown that RSSC IS are highly mobile under lab conditions and may contribute to host competitiveness and environmental stress tolerance (28). However, thus far our understanding of IS in RSSC is based on experimental studies of individual strains (27,28) and a small number of genomes derived from publicly available databases (29). As a result, the wider diversity and distribution of IS in RSSC is unknown. In addition, we poorly understand to what extent IS-driven variation follows the host phylogeny, concealing whether IS generally transmit vertically or if they are more often characterised by horizontal movement between lineages.

RSSC strains have broad host ranges infecting over 200 plant species within at least 50 families (30,31). They contain a bi-partite genome comprised of a chromosome and a megaplasmid, and are genetically diverse, being classified into four lineages, termed phylotypes (32). Phylotypes generally follow their geographical origin: Phylotype I includes strains originating primarily from Asia, Phylotype II from America, Phylotype III from Africa and surrounding islands in the Indian ocean, and Phylotype IV from Indonesia, Japan, and Australia (33). The four phylotypes have been redefined as three separate species, including *R. solanacearum sensu stricto* (Phylotype II), *R. pseudosolanacearum* (Phylotypes I and III) and an array of *R. syzygii* subspecies (Phylotype IV) (34). Considerable variation exists between and within *R. solanacearum* lineages regarding their metabolic versatility (35), tolerance to environmental stresses including starvation and low temperatures (36,37), and disease severity (38). This has been linked to a diverse accessory genome (39,40) which includes mobile genetic elements such as prophages (29,41). A recent analysis of RSSC prophages found that, while prophages were highly diverse, they were generally bacterial host phylotype-specific (41). In addition, prophage content tightly followed the host phylogeny with genetically similar hosts containing similar prophages. Some IS families have been reported to also have lineage-specific distributions (25), being exclusively found in specific RSSC species. Therefore, they may have similar distribution patterns to prophages and have limited horizontal transfer between compared to within bacterial lineages. Addressing this hypothesis requires a deeper exploration of the relationship between IS content and the host phylogeny and an assessment of potential IS fitness effects across the RSSC.

The large diversity of the strains within the complex and the economic losses associated with the disease make the RSSC a salient target for IS content analysis. While a recent study made a significant contribution to understanding RSSC IS using publicly available RSSC genomes (25), it had a sampling bias with low representation of strains from phylotypes IIA and IIB, missing a subset of hosts which are cold-adapted (36). In this study, IS were identified in a new representative collection of 356 RSSC isolates. These included isolates from all four phylotypes and six continents, with extensive sampling of phylotypes I and IIB. We specifically aimed to: i) characterise the total diversity and distribution of IS in RSSC, ii) assess the relationship between IS content and the host phylogeny; and iii) investigate the potential impact of IS movement on host fitness-associated genes. IS were initially identified in 27 Nanopore-assembled and five complete reference genomes with ISEScan (42) to determine IS diversity. Representative IS were then used to identify IS in all isolates using short read data with ISMapper (43). IS distributions were characterised by assessing lineage-specificity, with the relationship between IS content and host genetic background determined by comparing IS Bray-Curtis and host genetic dissimilarities. Finally, potential IS fitness effects were investigated based on their proximity to neighbouring genes.

## Methods

### RSSC hosts, sequencing, and genome assembly

RSSC hosts were selected from the National Collection of Plant Pathogenic Bacteria (NCPPB) and other reference strains maintained at Fera Science Ltd., York, UK. Genomic DNA extraction was performed on 384 isolates using Qiagen DNeasy Blood and Tissue Kit (DNeasy® Blood & Tissue Handbook, Qiagen, Hilden, Germany, 2020) followed by quantification of double stranded DNA product using Quantit dsDNA Assay Kit Broad range and Nanodrop (Thermo Fisher Scientific, Waltham, MA, USA). Host DNA was sequenced using Illumina MiSeq at the Earlham Institute, UK. 27 isolates were chosen for long read re-sequencing with Oxford Nanopore MinIon which was performed by the technological facility at the University of York. Guppy (https://nanoporetech.com/) was used for basecalling and hybrid assemblies were produced using the Unicycler (v0.4.8) pipeline on strict mode (44). After assembly some small contigs were filtered out based on sequence similarity and size. Short read sequence quality was assessed using FastQC (45) and trimming of adapters and low-quality ends was performed using Trimmomatic (v0.39) (46). Genomes were then assembled into draft assemblies using Unicycler (v0.4.8) on strict mode (44). To classify the genomes, a pangenome analysis was performed on 357 high quality genome assemblies plus 48 complete genomes downloaded from NCBI Genbank (Accessions available in Supplementary Data) and *R. picketti* 12b used as an outgroup. Core genome alignment was generated using Panaroo (v1.2.4) (47) with strict mode and MAFFT aligner (48). Phylogenetic tree was then constructed with IQ-TREE (49) and GTR+G4 model and monophyletic branches were assigned to phylotypes based on the close clustering with known phylotype (Figure S1; Table S1).

### IS detection in Nanopore and reference genomes

IS were detected in 27 Nanopore-assembled genomes (Table S2) and five complete reference genomes (GMI1000, phylotype I; K60, phylotype IIA; UY031, phylotype IIB; CMR15, phylotype III; PSI07, phylotype IV), representing different phylotype lineages in the RSSC phylogeny. Firstly, putative full-length IS were identified with ISEScan v.1.7.2.3 (42) using the removeShortIS parameter to remove partial IS copies. In total, 2,768 full length IS were identified. These were filtered by removing IS duplicates based on whether they belonged to the same IS family and had the same length or if they had identical terminal inverted repeats. After filtering, 861 IS were retained and used for subsequent analyses. IS diversity was then assessed by determining IS sequence similarity using Mash v2.2 (50) and generating a pairwise Mash distance matrix using the “mash triangle” function with sketch size = 10,000. IS were clustered based on sequence similarity using K-means clustering with the R package ‘pheatmap’ v1.0.12. The optimal number of clusters was determined using a Silhouette plot with the R package ‘factoextra’ v1.0.7. A total of 66 IS clusters were identified and one IS representative was selected at random from each cluster. To determine IS identities, IS representatives were blasted against the ISFinder database (https://isfinder.biotoul.fr/) with successful hits determined using an E-value < E^-50^ threshold. For successful hits, the ISFinder database copy of the IS was downloaded.

### IS detection in RSSC isolates from short read data

IS were identified in all 356 RSSC isolates using only short read data with ISMapper v.2.0.2 (43). Briefly, ISMapper maps short read data to reference IS, identifying reads that map to and overhang the 3’ and 5’ IS flanks. Mapped reads are then further mapped to an annotated reference bacterial genome and IS positions are determined where 3’ and 5’ flanking reads both map to similar genomic locations. Representative IS sequences downloaded from the ISFinder database were used as references. As gene content varies between RSSC lineages, different reference bacterial genomes were used depending on the lineage of the isolate being analysed. As a result, depending on their phylotype classification, reads from isolates were mapped to GMI1000, K60, UY031, CMR15, and PSI07 strains. Annotated bacterial genomes (GenBank format file, gbff) were downloaded from the NCBI database (Table S3). After running ISMapper, IS hits were filtered to remove potential false positives. IS with unknown 5’ or 3’ coordinates were removed. Further, IS hits that overlapped within the same isolates were de-duplicated as they likely represent different reference IS mapping to the same location. As all overlaps occurred with IS from the same family, the remaining de-deuplicated IS were given an “Unknown_” + IS family label (e.g Unknown_IS5). Finally, IS that were found to disrupt transposases, likely representing intra-IS insertions, were removed. This is because IS that map inside multi-copy genes will map to all copies irrespective of the true IS content and therefore may generate spurious hits (43). To see whether IS detection with short read data has lower accuracy than with long read assemblies, IS copy number from each method was compared using the 27 Nanopore-sequenced strains. While there was a significant correlation in IS copy number when using short and long read data (Kendall’s p < 0.01; Figure S2A), short read data had lower IS detection power, identifying on average only ∼73% of the IS found with long read assemblies. Therefore, to fairly compare IS between all strains, IS were only detected with short read data. Moreover, as read depth can affect IS detection power, the average per base read depth for each isolate was determined and compared between host phylotype lineages (Figure S2B). Read depth was calculated by first aligning paired-end reads to the isolate chromosome or megaplasmid using the Burrows-Wheeler aligner (51) “bwa mem” command. Per base read depth was then determined using SAMtools (52) “sort” and “depth” commands, including bases with no coverage. Although phylotype IIB strains had significantly greater read depth than phylotype I (Kruskal-Wallis: x^2^ = 75.1; d.f = 4, p < 0.01), there was no significant difference in read depth for all other pairwise comparisons. Further, as ∼99% of isolates from each phylotype (except for IIA with 93%) had an average per base read depth > 30x (average read depth: chromosome = 67.9x ± 17 s.d; megaplasmid = 59.9x ± 14.2 s.d), which is above the suggested threshold for correct IS detection (43), read depth likely had a minimal impact on IS detection and copy number.

### Determining the relationship between IS content and host genetic background

The relationship between IS content and host genetic background was first assessed by comparing IS profiles between host lineages using principal coordinate analysis. Differences in IS subgroup content between isolates were determined by calculating IS Bray-Curtis dissimilarities with the R package ‘vegan’ v.2.5-7, which accounts for presence, absence, and relative abundance of IS subgroups in host genomes. Principal coordinate analysis of Bray-Curtis dissimilarities was conducted using the R package ‘ape’ v.5.6-1 (53) and IS content differences between phylotypes were tested statistically using ANOSIM with 9,999 permutations.

IS content and bacterial host genetic background was further compared by calculating the congruence between the RSSC phylogeny and a UPGMA tree constructed using IS Bray-Curtis dissimilarities. The IS Bray-Curtis UPGMA tree was constructed from a pairwise Bray-Curtis dissimilarity matrix using the R package ‘phangorn’ v.2.8.1 (54). A tanglegram between the RSSC ML tree and the IS Bray-Curtis UPGMA tree was generated with functions in the R package ‘ape’, using the R package ‘phytools’ v.1.0-1 (55) to rotate the RSSC ML tree to minimise connected lines crossing between the trees. Congruence between the RSSC ML tree and the IS Bray-Curtis UPGMA tree was assessed using Procrustes Approach to Cophylogenetic Analysis (PACo) v.0.4.2 (56) in R. Briefly, cophenetic distance matrices were constructed using the IS Bray-Curtis and RSSC phylogenetic trees. The distance matrices were then transformed into principal coordinates and compared with each other using a Procrustean superimposition with a null model (host phylogenetic tree ordination does not predict IS UPGMA tree ordination, i.e., there is no congruence) to determine tree congruence. Statistical significance was calculated based on 1,000 network randomizations under the “r0” randomization model.

#### Data visualisation and statistical analysis

Statistical analyses and data visualisation were carried out using Microsoft Excel v.2102, R v.4.0.3 and RStudio v1.4.1103. The difference in IS copy number between short read IS detection (ISMapper) and long read detection (ISEScan) was determined using a Kendall-rank correlation. Read depth and IS copy number was compared between phylotypes using Kruskal-Wallis tests followed by Dunn’s post-hoc test. The difference in IS copy number and the number of IS close to (< 100 bp from start codon) or inside genes in the chromosome and megaplasmid were tested using paired t-tests. Graphs and heatmaps were made using the R package ‘ggplot2’ v.3.3.3. The Bray-Curtis UPGMA and ML phylogenetic trees were visualised using the R ‘ggtree’ package v.2.1.4 (57).

## Results

### Insertion sequence abundance and distribution across *Ralstonia solanacearum* species complex

We first compared insertion sequence content and distribution across the genomes of 356 RSSC bacterial isolates. All bacterial isolates contained IS and a total of 24,732 IS were identified. A significantly higher number of IS were identified in the megaplasmid (12,734 IS) than in the chromosome (11,998 IS; t = 9.52; d.f. = 355; p < 0.001, Figure S3) despite the megaplasmid being approximately half the size of the chromosome (32). Insertion sequences belonged to eleven IS families that included IS5, IS3, IS110, IS256, IS21, IS701, ISL3, IS4, IS630, IS1595, and IS1182. The most prevalent IS families were IS5 (78.9%) and IS3 (15.4%), with the remaining families each comprising less than 2% of the total number of IS (Figure 1A). However, greater IS prevalence in the megaplasmid was not consistent across all IS families and IS from IS110 and IS3 families were predominantly detected in the chromosome. IS copy number was significantly different between phylotypes overall (Kruskal-Wallis: x^2^ = 151.1; d.f = 4, p < 0.001). However, this result was driven by uneven representation of isolates across phylotypes (I (59), IIA (15), IIB (269), III (8), and IV (5)), which resulted in overrepresentation of two insertion sequences that were common to phylotype IIB bacterial isolates: IS5 and IS3 (Figure 1C). Nonetheless, IS5 and IS3 were highly prevalent across the RSSC phylogeny, despite the oversampling of phylotype IIB isolates (Figure 1B, C). In addition to IS5 and IS3, certain lower abundance IS families, such as IS110 and IS256, were also present across the phylogeny. However, a small number of families were found in specific lineages. For example, phylotype I isolates almost uniquely contained ISL3 and IS4 and most of the IS701 (89.6%) and IS1595 (72%) IS. In addition, phylotype IIB exclusively contained IS21 IS. Therefore, whilst RSSC IS family content was dominated by IS5 and IS3, lineage-specific presence-absence patterns were observed across the phylogeny.

**Figure 1.**
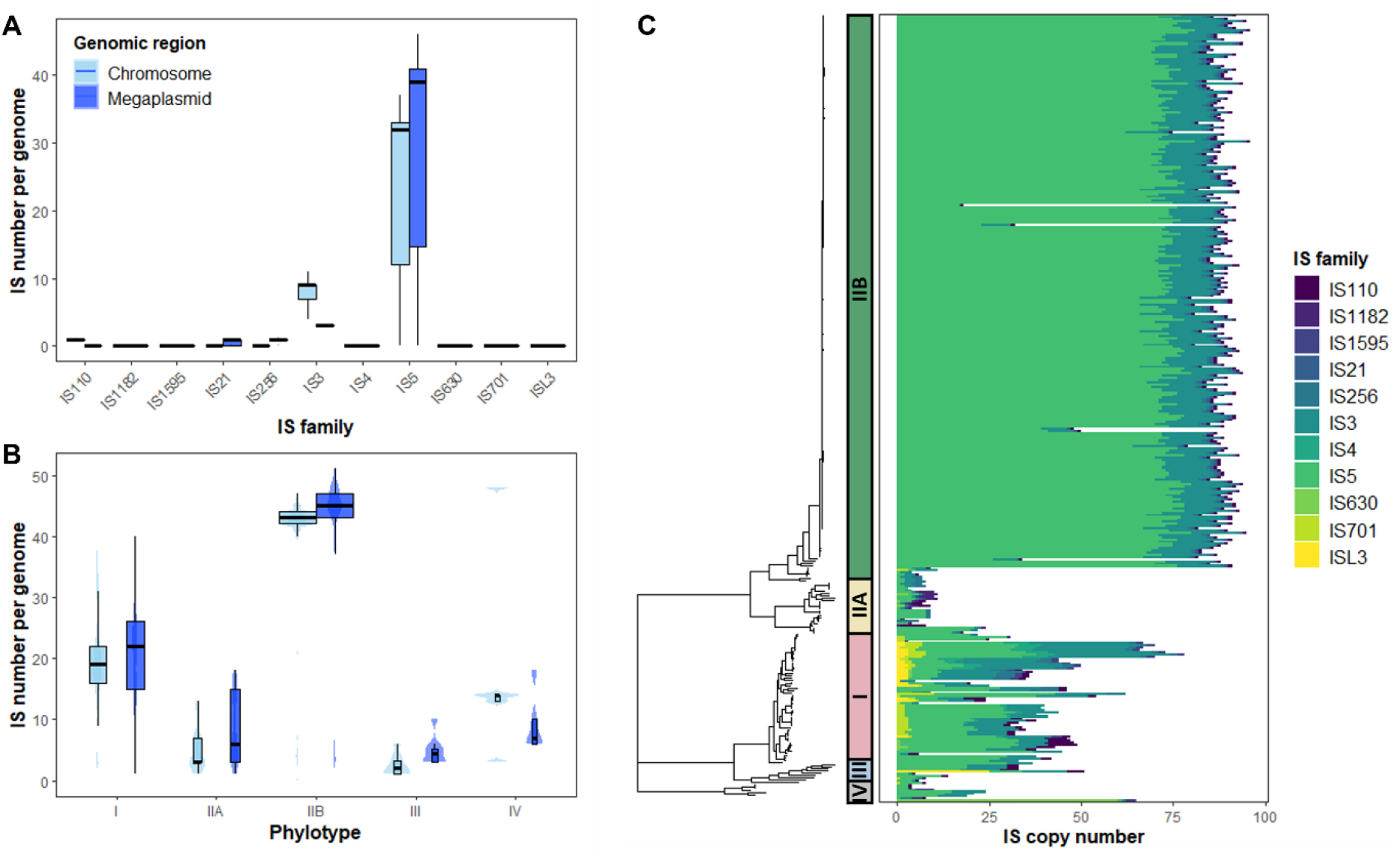
IS number and content varies across the RSSC phylogeny. A) Box plot and violin plot of IS prevalence for each IS family for both the chromosome and megaplasmid. B) Box plot and violin plot showing the number of IS in isolates from each lineage in the phylogeny. For both A and B, IS prevalence in both the chromosome (light blue) and the megaplasmid (dark blue) is shown. C) Left side is RSSC phylogeny with coloured bars showing phylotype label, right side is a heatmap of IS prevalence coloured by IS family

#### Insertion sequence content tracks host phylogeny at IS subgroup level

IS family distributions were further investigated by looking at the distributions of intra-family IS subgroups (Figure 2A). While the highly prevalent IS5 and IS3 families were found to contain eight and nine IS subgroups, respectively, only individual IS subgroups were identified for other IS families. In contrast to IS5 and IS3 families that were generally widespread and found in all lineages, IS5 and IS3 subgroups showed lineage-specific distributions. The IS5 subgroup IS1021 comprised most of the IS5 IS in the large clonal IIB sub-lineage and was present only in low abundance in other lineages. The remaining IS5 subgroups were primarily found in phylotype I isolates, although some were also found in phylotypes IIA and IV. Similarly, IS3 subgroups in the clonal IIB sub-lineage primarily included ISRso10 and ISRso20, whereas the remaining IS3 subgroups were mainly found in phylotype I. IS subgroup lineage-specificity was further verified using a principle-coordinate analysis based on host IS subgroup Bray-Curtis dissimilarities (Figure S4). Phylotypes could be significantly distinguished based on their IS contents (ANOSIM: p < 0.001 for all lineages), with particularly strong clustering between the clonal IIB sub-lineage and phylotype I isolates. Notably, a more diverse cluster was also observed containing isolates from all phylotypes, including non-clonal IIB isolates and all IIA, III, and IV isolates. Although overlaps were small with phylotype III isolates, the general clustering between these lineages suggests that their IS contents were similar. Together, these findings suggest that intra-family IS subgroups were mainly phylotype-specific.

**Figure 2.**
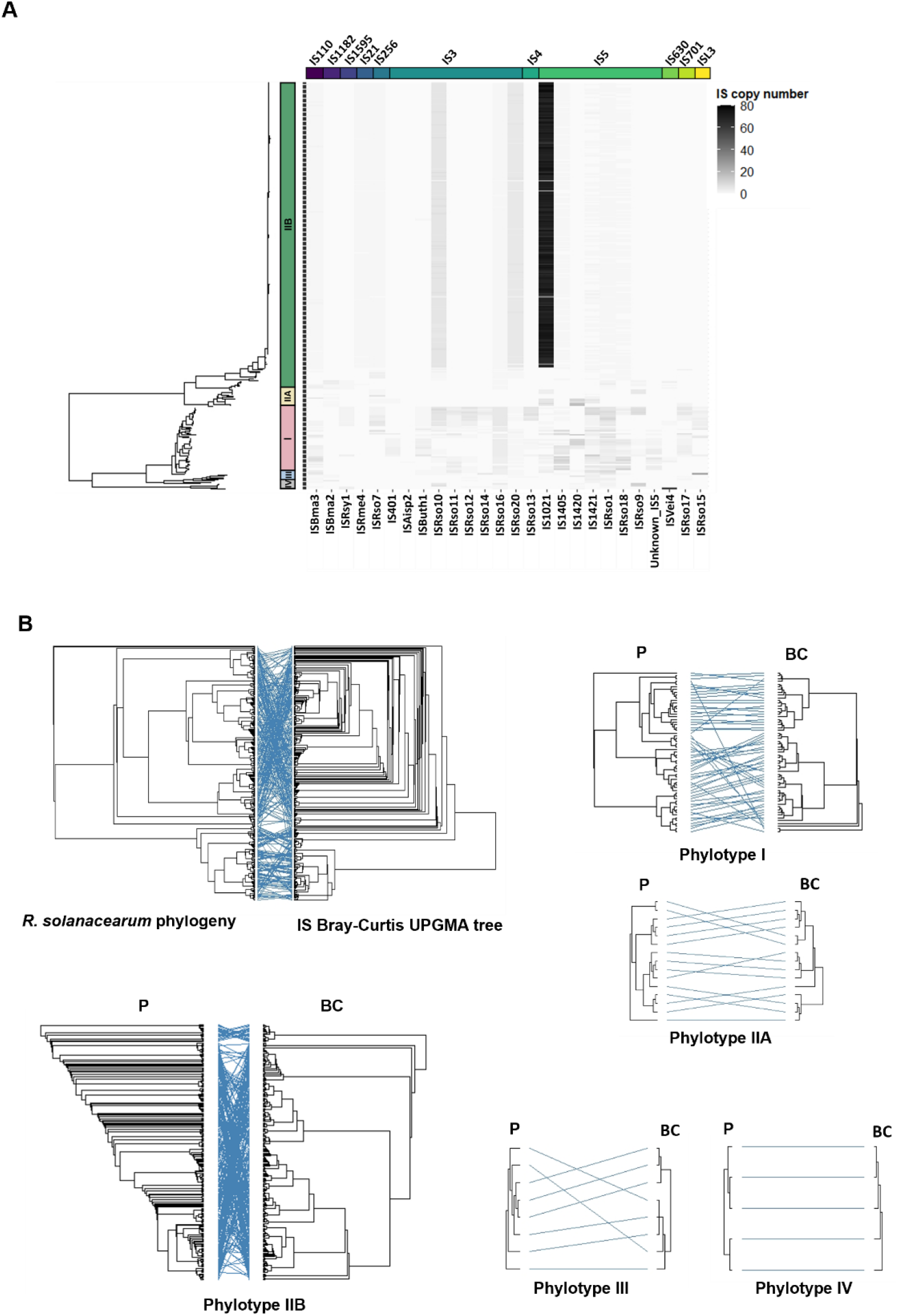
IS subgroups generally show host lineage-specificity and widespread IS are found in different genomic regions depending on host lineage. A) Heatmap of IS subgroup prevalence across the RSSC phylogeny. IS subgroups clustered by IS family, shown with coloured bars above the heatmap. B) Tanglegrams showing the congruence between RSSC phylogeny (left side) and UPGMA tree (right side) calculated using Bray-Curtis dissimilarity of IS presence. The first tanglegram shows congruence for the whole phylogeny and the other plots show congruence within lineages. Phylogeny is labelled with P and UPGMA tree is labelled with BC (Bray-Curtis).

To test if the observed host lineage-specificity of IS subgroups could be explained by host genetic similarity, we measured the congruence between the RSSC phylogeny and a UPGMA tree constructed based on IS subgroup Bray-Curtis dissimilarities (Figure 2B). Significant congruence was detected between the total RSSC phylogeny and Bray-Curtis UPGMA tree (M^2^_xy_ = 0.34, p < 0.001, N = 1,000). To ensure this wasn’t biased by the large clonal IIB sub-lineage, the analysis was repeated for each lineage independently, including phylotypes III and IV which had relatively lower sampling sizes. Except for the small sampling size phylotype III, significant congruence was found when analysing lineages independently (phylotype I - M^2^_xy_ < 0.001, p < 0.001, N = 1,000; IIA - M^2^_xy_ < 0.001, p < 0.001, N = 1,000; IIB - M^2^_xy_ = 0.005, p < 0.001, N = 1,000; III - M^2^_xy_ < 0.001, p = 0.052, N = 1,000; IV - M^2^_xy_ < 0.001, p < 0.05, N = 100) and each lineage-level congruence comparison had improved goodness-of-fit compared to the congruence analysed at the level of the whole strain collection. This analysis therefore confirms that genetically similar hosts contain similar IS contents.

#### Insertion sequence subgroups are found in different genomic regions in different host lineages

While IS subgroups were generally lineage-specific, they were often found in low abundance in other lineages, indicative of recent horizontal IS gain events rather than vertical transmission from a common ancestor. This was investigated by comparing the prevalence of IS in the chromosome and megaplasmid between lineages (Figure S5). Some IS subgroups were found in the same genomic region in all lineages. For example, the IS3 subgroup ISRso10 was mainly inserted into the chromosome, and the IS5 subgroups IS1021, IS1405, IS1421, and ISRso18 were primarily found in the megaplasmid. However, 42% of IS subgroups (11/26) had different genomic locations that depended on the host lineage. For example, the IS1182 subgroup ISBma2 was found in the chromosome in phylotype IIB and III strains but was only found in the megaplasmid in phylotype IIA. In addition, IS subgroup genomic region was not conserved within IS families; in phylotype IIB isolates, the IS3 subgroups ISRso10 and ISRso20 were found in the chromosome and megaplasmid, respectively. In contrast, especially in phylotype IIB, IS5 subgroups were primarily found in the megaplasmid. These results suggest that, irrespective of their family background, IS subgroups are found in different genomic regions within different bacterial host lineages.

#### Insertion sequences potentially disrupt genes associated with virulence and competitiveness and may contribute to inter-phylotype trait variation

In the RSSC, IS insertions have been shown to occur near to or inside of type III effectors and global virulence regulators, affecting host virulence and phenotypic plasticity (27,29). Therefore, the proximity of IS to neighbouring genes was investigated. The distance of IS to their closest neighbouring gene was found to have a roughly exponential distribution, with most IS being physically close to coding sequences (Figure 3, inset). Indeed, 71.4% of all IS identified were found less than 100bp from a neighbouring gene’s start codon, including 78.7% of chromosomal IS and 66.4% of megaplasmid IS. Notably, although coding sequences comprise approximately 86.8% of the RSSC genome (32), only 4.9% of IS were predicted to disrupt genes (7.1% of chromosomal IS and 2.8% of megaplasmid IS) potentially changing their function. Despite IS being significantly more prevalent in the megaplasmid than the chromosome, isolates contained significantly fewer gene-proximate (closer than < 100 bp to a start codon) and intra-genic IS in the megaplasmid than in the chromosome (t = -15.60; d.f. = 353; p < 0.001, Figure S6). Moreover, IS gene disruptions were not caused by all IS subgroups (Figure S6), and subgroups that mainly inserted into inter-genic regions were found in most IS families, including IS5 and IS3, which suggests that potential gene disruptions were not IS family-specific. However, while some IS subgroups caused gene disruptions more often than others, the proportion of IS insertions within genes differed between lineages (Figure S7). Five IS subgroups, including ISAisp2, ISRso14, ISRso16, ISRso13, and ISRso9 caused > 90% of their gene disruptions in individual lineages, and only eight IS subgroups caused > 10% IS disruptions in more than two lineages.

**Figure 3.**
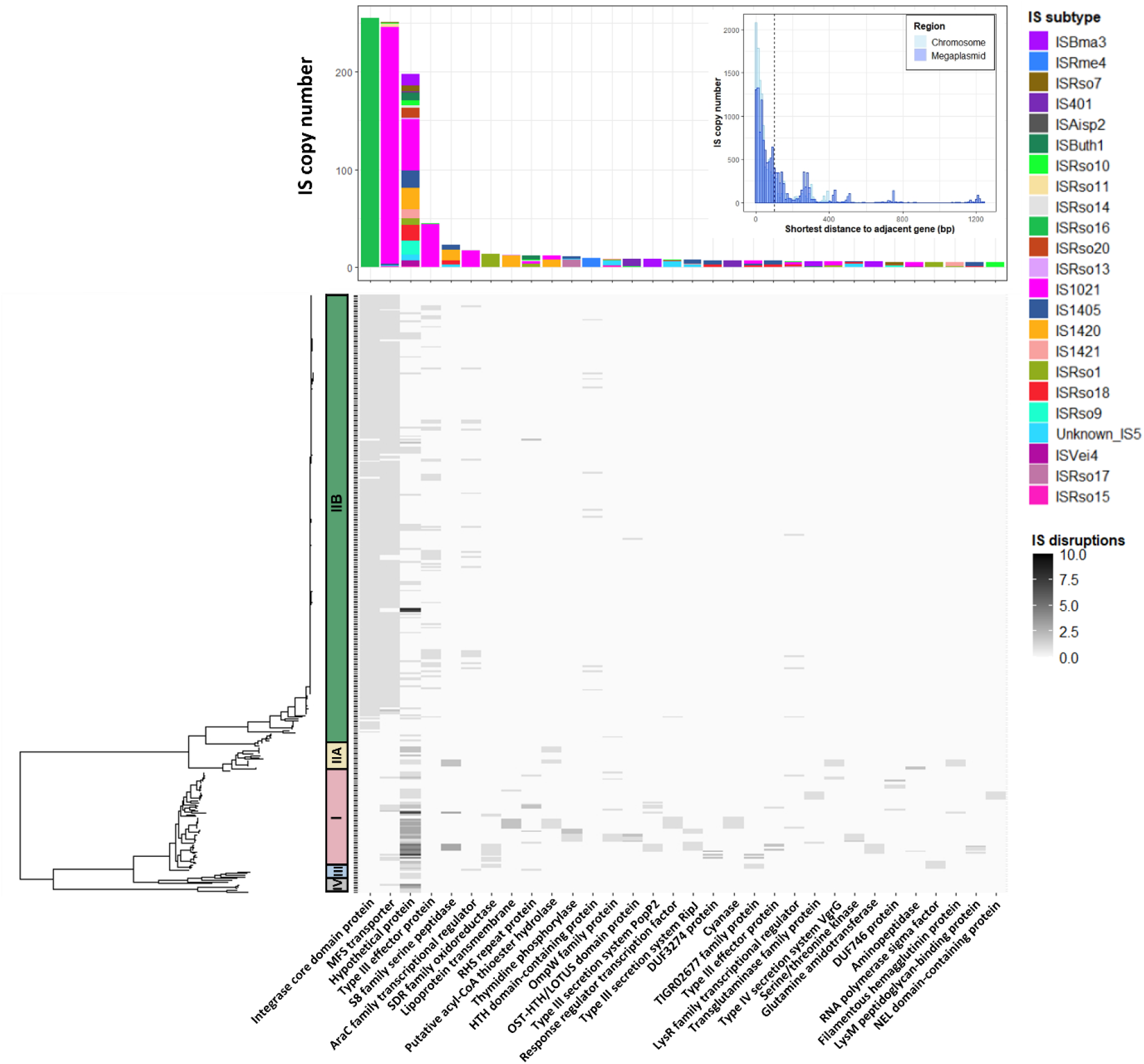
IS disrupt genes across the phylogeny in a lineage-specific manner. Inset) Histogram of the shortest predicted distance between each IS and its adjacent gene’s start codon. Histograms for chromosomal and megaplasmid IS are overlaid and coloured separately. Dotted vertical line shows 100 bp threshold. Bottom; heatmap showing the number of IS-mediated disruptions in different gene types across the phylogeny. Only genes that contained disruptions in five or more isolates are shown. Genes are ordered by frequency of IS disruption. Top; stacked bar chart showing the IS subgroup distribution for each gene disrupted.

In total, IS were found to disrupt 335 genes across all isolates, including 38 genes that were disrupted multiple times in separate loci. This excludes hypothetical proteins, whose functions remain unknown, which were disrupted in 95 loci. Genes that were disrupted by IS in more than five isolates were analysed further (Figure 3). Of these, 32 disrupted genes were identified and associated with potential fitness functions, including bacterial virulence, antibiotic and oxidative stress tolerance, protein modification, cellular metabolism, and RNA-binding. Most gene disruptions (20/32) occurred in closely related isolates in phylotype I, indicating they could have arisen via vertical transmission from a common ancestor. However, some gene disruptions also occurred in a small number of more distantly related isolates while others, such as those in S8 family serine peptidase, RHS repeat protein, and LysR transcriptional regulator, were mainly found in distantly related isolates across multiple lineages. Except for IS disruptions in the clonal IIB sub-lineage, widespread gene disruptions were generally caused by multiple IS subgroups in different isolates, suggesting they likely represent parallel insertions. IS disruption of these genes may be under stronger selection and provide a fitness benefit to their hosts.

## Discussion

Insertion sequences contribute to genetic diversity in bacterial pathogens, affecting a myriad of traits including metabolism (10,58), virulence (7,8,59), and stress tolerance (4,6). In this study, we analysed insertion sequence content in the plant pathogenic RSSC bacterium using a diverse, global collection of 356 RSSC strains. IS were identified in all strains and belonged to eleven IS families, the most prevalent being from the IS5 and IS3 families. Although IS families tended to be widespread, IS subgroups were host lineage-specific and genetically similar hosts had similar IS contents. IS subgroups were rarely found in multiple lineages and, when found, were inserted into different genomic regions with different lineages, indicative of potential horizontal movement between lineages. IS were generally close to neighbouring genes and caused disruptions in several genes, potentially associated with bacterial virulence and stress tolerance. Overall, our results suggest that IS elements might be evolving in tandem with their hosts, potentially contributing to the phenotypic diversity and fitness of the RSSC.

A recent analysis of RSSC insertion sequences provided important insights into the potential diversity and distribution of IS across the RSSC (25). Here we built on this analysis by including a more representative sampling of new RSSC genomes, including isolates from IIA and IIB phylotype groups that were underrepresented in the previous study. Overall, our results support the previous findings (25). IS were identified in all isolates analysed although IS copy number varied across the RSSC. We found that a large clonal phylotype IIB sub-lineage contained the greatest number of IS, followed by phylotype I and IV, with IIA, III, and non-clonal IIB isolates containing very few IS. Excluding the clonal IIB sub-lineage, our results support previous work suggesting phylotype I has the highest IS copy number (25). Further, the most prevalent IS were from the IS5 and IS3 families, which were widespread across the RSSC with high copy number in the clonal IIB sub-lineage and in phylotype I isolates. Therefore, these IS families are likely the dominant IS families in the RSSC. However, in contrast with previous findings, non-clonal IIB isolates had very low IS copy number, and across all lineages, we detected fewer IS copies per isolate than were found previously. This is likely due to methodological differences as, while Goncalves *et al* (25) identified IS in complete genome sequences, we identified IS with short read data which had lower IS detection power than with long read assemblies. This could be because IS detection with short reads depends on reference IS identified from long read assemblies and we identified fewer reference IS than were found previously. In addition, we filtered out intra-IS insertions to avoid spurious hits which, depending on the prevalence of intra-IS insertions across the RSSC, may have resulted in under-estimated IS copy numbers.

Consistent with the findings of Goncalves *et al* (25), we found that most IS families (6/11) that were less abundant were also less widespread: ISL3, IS701, IS4, IS630, IS1595, and IS21 IS had low prevalence (collectively 2.7% of total IS) and were mainly found in specific lineages (phylotype I: ISL3, IS701, IS4, IS1595; phylotype IIB, IS21; phylotype IV, IS630). However, an important extension in our analysis included assessing the distribution of intra-family IS subgroups, enabling the detection of nuanced patterns that sometimes occur within IS families (60). We found that widespread IS5 and IS3 families contain many IS subgroups which have strong lineage-specific distributions. Indeed, host lineages were significantly distinguishable based on IS subgroup content alone. Consequently, IS subgroups appear to be largely restricted within distinct host lineages. Differential IS family copy numbers between lineages have previously been detected in other pathogens (61) and observed to rise in long-term experimental studies (62). Whilst it is unclear whether these IS families included single or multiple IS subgroups, our results suggest that IS subgroups should be considered in future analyses as their distributions may differ between lineages that contain similar IS family copy numbers. The relationship between IS content and host genetic background was further analysed by calculating the congruence between the host phylogenetic and a UPGMA tree constructed based on IS subgroup Bray-Curtis dissimilarities. We found that there was significant congruence between the trees overall and independently within lineages, suggesting that genetically similar hosts contain similar IS. These findings mirror those of a recent study conducted on RSSC prophages (41) which found similar lineage-specific distributions and detected significant congruence between prophage content and the host phylogenetic tree. Therefore, multiple mobile genetic elements in RSSC appear to track the evolution of their hosts and act as sources of genetic diversity between lineages.

In contrast with previous analyses (25), we found significantly more IS in the megaplasmid than the chromosome. This was surprising given the RSSC megaplasmid is approximately half the size of the chromosome (32). One potential explanation is that, although the chromosome and megaplasmid have a similar number of coding sequences relative to their size, over 90% of the RSSC core genome is found in the chromosome (32,63). This includes most house-keeping genes (32) whose disruption would likely reduce host fitness. Therefore, given the megaplasmid mainly contains accessory genes with non-essential functions, it may be under weaker purifying selection against IS insertions and so undergoes greater IS propagation. Notably, greater megaplasmid IS prevalence was not found for all IS families and the IS110 and IS3 families were mainly found in the chromosome, likely due to their chromosomal prevalence in the large clonal IIB sub-lineage. Although many IS have little to no target specificity (64), IS distributions were inconsistent with random insertions into the chromosome and megaplasmid; if IS insertions were random then the megaplasmid should have proportionately fewer insertions compared to the chromosome in all lineages due to its smaller size (32), which was not the case. This could reflect lineage-specific purifying selection against IS inserting into specific genomic regions. Alternatively, some IS are highly specific and target sites of DNA replication (65,66), motifs upstream of promoters (67,68), and secondary DNA structures (69–71). Therefore, IS may have had target preferences in the chromosome or megaplasmid within each lineage if insertion targets vary phylogenetically. These hypotheses should be addressed in future studies through analyses of RSSC IS target sites and IS fitness effects. The distribution of IS between the chromosome and megaplasmid was further investigated by comparing the prevalence of IS subgroups in each genomic region between lineages. Interestingly, although some IS subgroups inserted into the same genomic region across the RSSC, most IS subgroups inserted into different regions depending on the host lineage. As IS subgroups tended to have high abundances in specific lineages and only low abundances in others, this may have arisen through inter-lineage horizontal IS transfer. IS movement between bacterial lineages has previously been attributed to plasmids (72,73), integrative and conjugative elements (74), and, rarely, prophages (73). Both integrative and conjugative elements and prophages have both been found to be lineage-specific (40,41) and therefore may have limited movement between lineages. However, RSSC strains are naturally competent and is capable of being transformed with DNA from different lineages (75–77). Therefore, IS horizontal transmission may occur via plasmids although additional analyses are required to investigate this.

Insertions nearby or inside genes can result in both gene disruption (5,58,78,79) and gene activation (80–82). In RSSC, IS insertions have been found to disrupt type III effectors (25) and global virulence regulators (27). We found that the majority of IS (71.4%) were inserted within 100bp of their closest neighbouring gene’s start codon and could potentially have affected gene expression (although the positions of IS relative to gene promoters is unclear). Such potential IS-associated fitness effects should be addressed further by analysing gene promoter positions in both the bacterial genomes and in the IS to determine whether promoters are disrupted or modified. Only 4.9% of IS were found to disrupt genes despite putative coding sequences comprising approximately 86.8% of the RSSC genome (32). Similarly, low proportions of IS-mediated gene disruptions have been found in other pathogens (61), suggesting there may be strong purifying selection against intra-genic insertions. Consistent with previous analyses (25), IS disruptions were found in various genes with potential roles in bacterial virulence, competition and stress tolerance, including type III effectors, membrane transporters, virulence regulators, and proteins involved in antibiotic and oxidative stress tolerance. Most gene disruptions were found in small genetically similar clades in phylotype I suggesting they have likely spread through vertical transmission and may provide local adaptive benefits. However, some disruptions were more widespread, and either found in more genetically distant isolates or spread across different lineages. Except for phylotype IIB disruptions, which were mainly caused by IS5 subgroup IS1021, these disruptions were generally caused by different IS subgroups in different isolates suggesting they might provide “general” fitness benefits, which are not dependent on the local environment.

In conclusion, this study provides insights into the distribution and potential spread of IS across the RSSC. Our results highlight that although IS are widespread on the family level, they are lineage-specific on the subgroup level and are tightly bound to the evolution of their hosts. Further, while IS generally cause lineage-specific gene disruptions, some genes are disrupted in multiple lineages by different IS subgroups. Therefore, IS may affect host fitness both within lineages and across the whole RSSC.

## Supplementary figures

**Figure S1.**
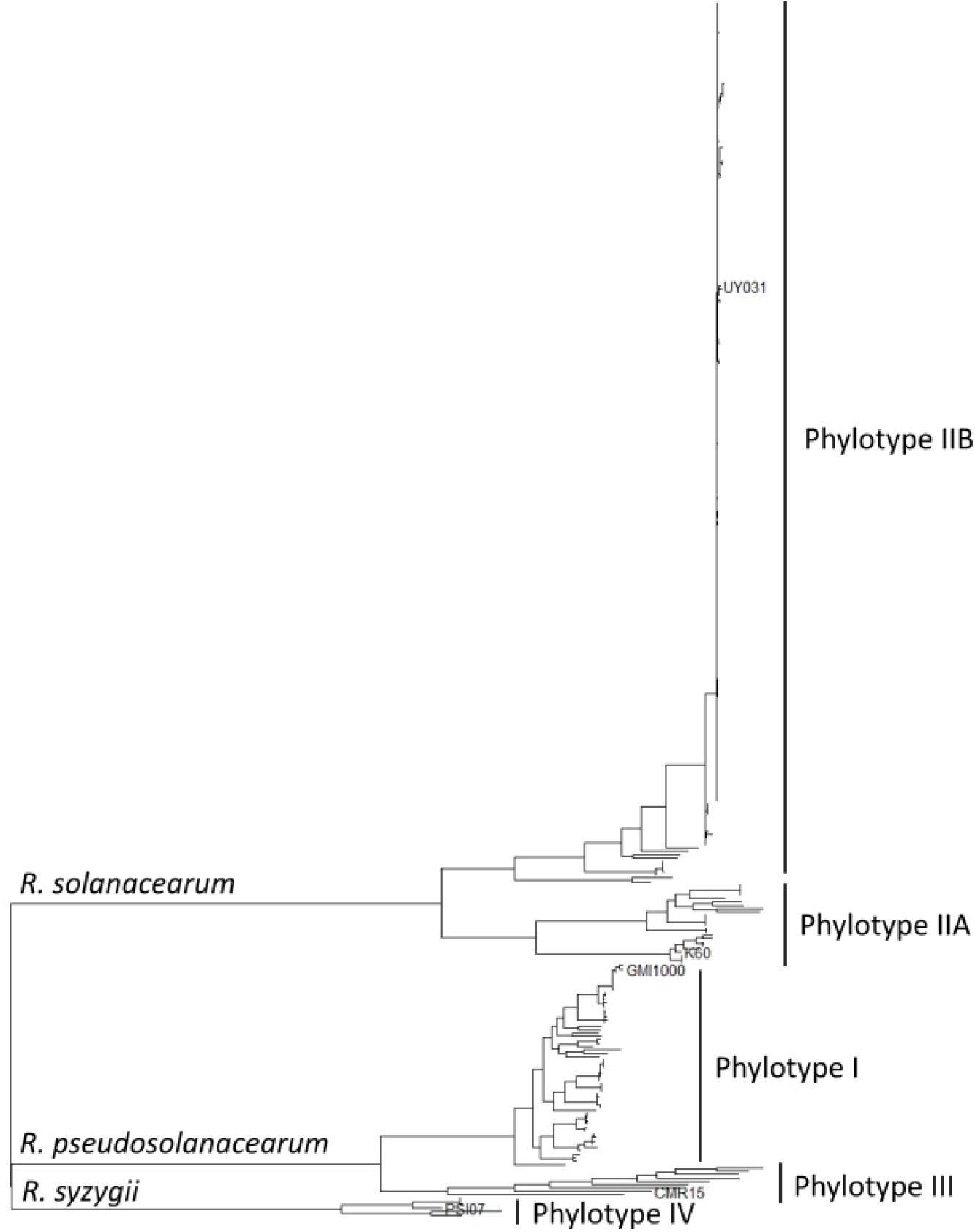
Phylogeny of *Ralstonia solanacearum* species complex. Maximum Likelihood phylogeny was constructed based on the genomes of 356 *Ralstonia solanacearum* species complex strains the National Collection of Plant Pathogenic Bacteria (NCPPB) and other reference strains maintained at Fera Science Ltd, along with 5 previously phylotyped and sequenced strains from NCBI Genbank (names shown at the tips of tree). Phylogenetic relationships between known phylotypes were used to assign the 356 strains sequenced in this study to given phylotype clusters

**Figure S2.**
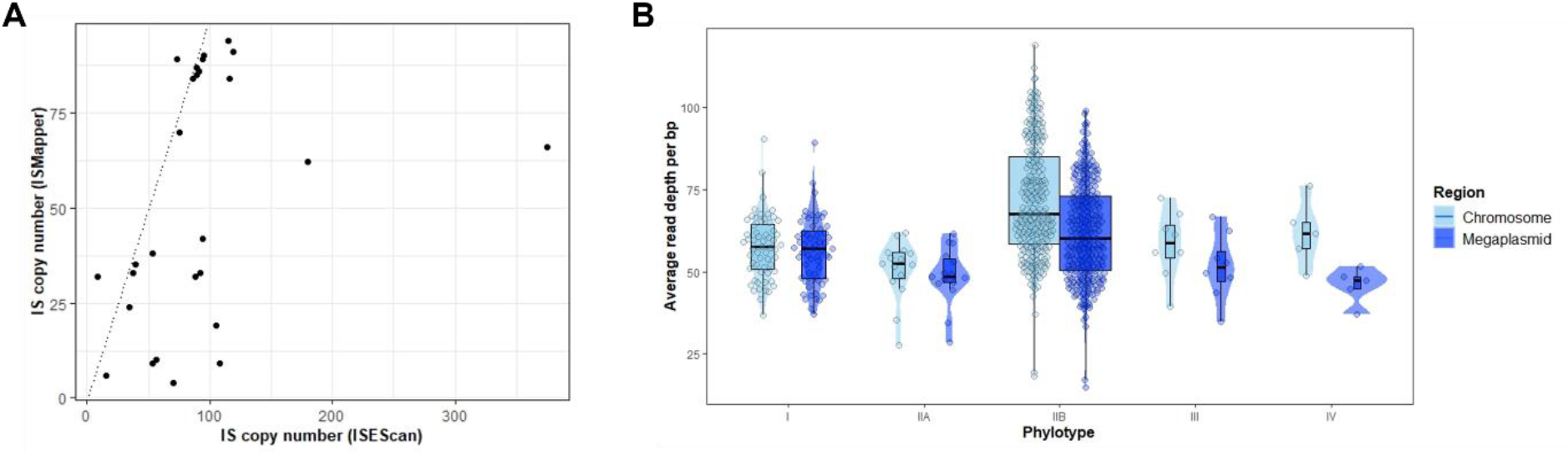
Short read IS detection is correlated with long read assemblies and is unaffected by average read depth. A) Scatterplot showing IS copy number determined using ISEScan (long read assemblies) against ISMapper (short read data). Dotted line has slope = 1, intercept = 0 and shows the expected relationship if both methods find the same copy number. B) Boxplot and violin plot showing average read depth per isolate for each phylotype. Data is scattered to reduce overplotting. The difference in read depth between phylotypes was tested statistically using a Kruskal-Wallis test.

**Figure S3.**
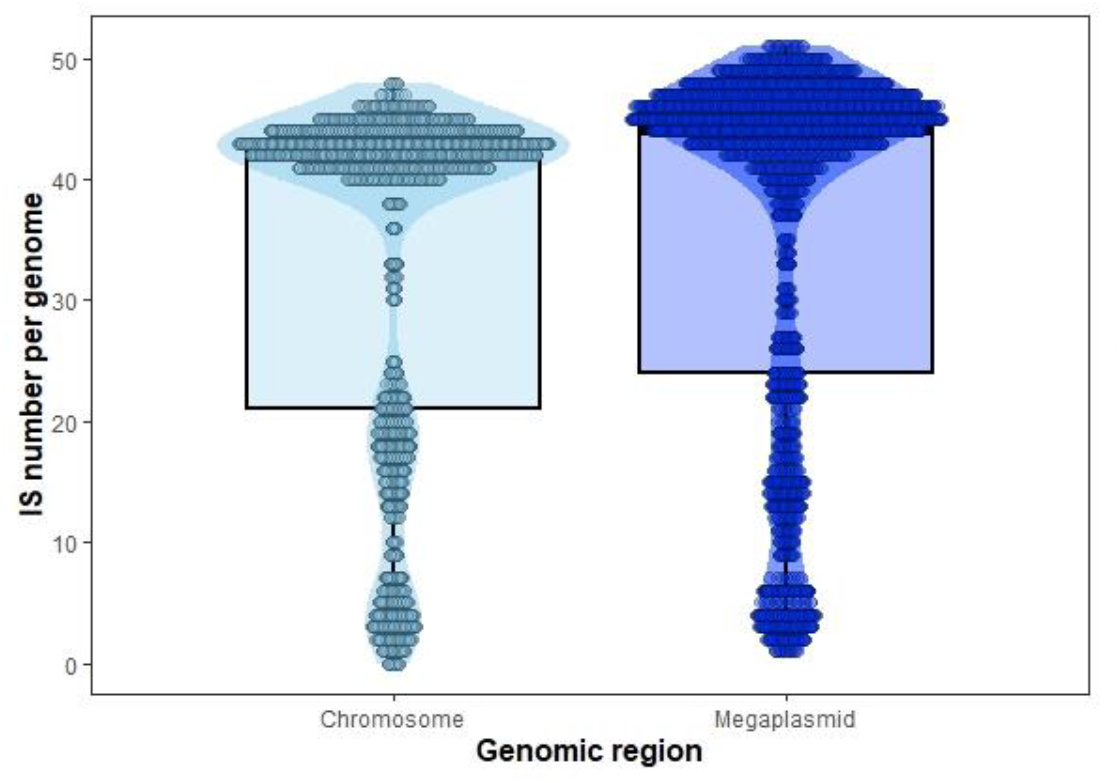
The megaplasmid contains significantly more IS than the chromosome. Boxplot showing IS copy number in the chromosome and the megaplasmid. Boxplots are coloured by genomic region. The difference in IS number between regions was tested statistically using a paired t-test.

**Figure S4.**
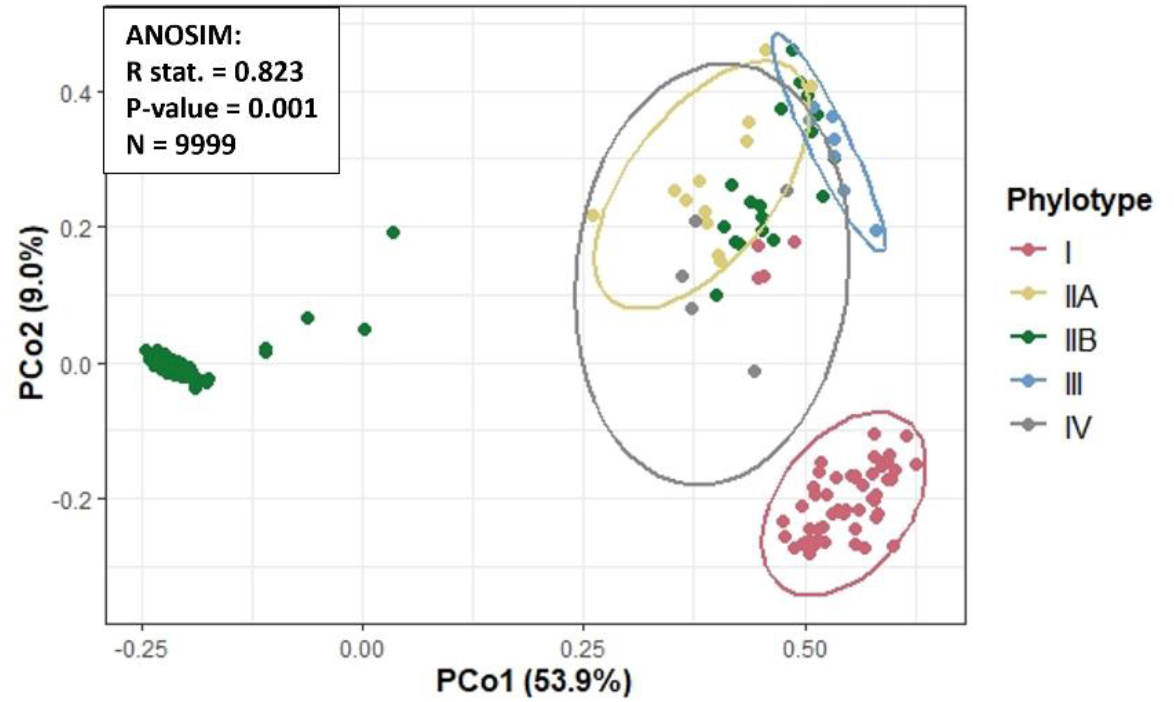
RSSC phylotype lineages have unique IS contents. PCoA plot based on pairwise isolate IS content Bray-Curtis dissimilarities. Points are coloured by host phylotype and an ellipse is plotted showing predicted phylotype point distributions. Phylotype IIB does not show an ellipse due to bimodal distribution caused by clonal and non-clonal sub-lineages. Box shows the results of ANOSIM including the R statistic (measure of phylotype IS distinguishability), P-value, and number of permutations.

**Figure S5.**
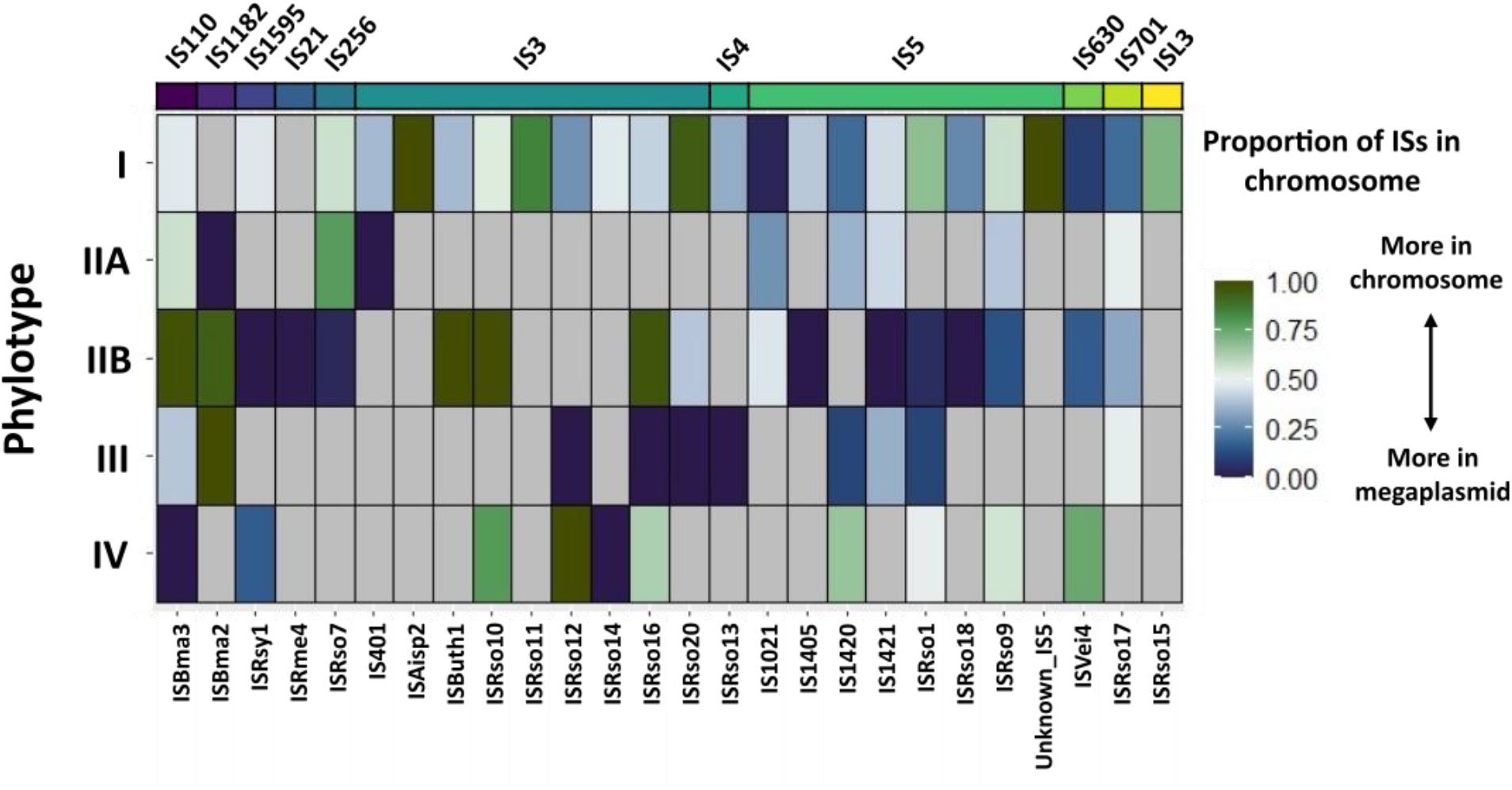
IS subgroups are located in different genomic regions in different lineages. Heatmap showing the proportion of each IS subgroup found in the chromosome in each host lineage (IS subgroup copies in chromosome/total IS subgroup copies for each lineage). Cells with a high proportion of chromosomal IS are shown in green and cells with a high proportion of megaplasmid IS are shown in blue. Grey cells are where an IS subgroup was not present in the lineage. IS are clustered in same order as Figure 2A.

**Figure S6.**
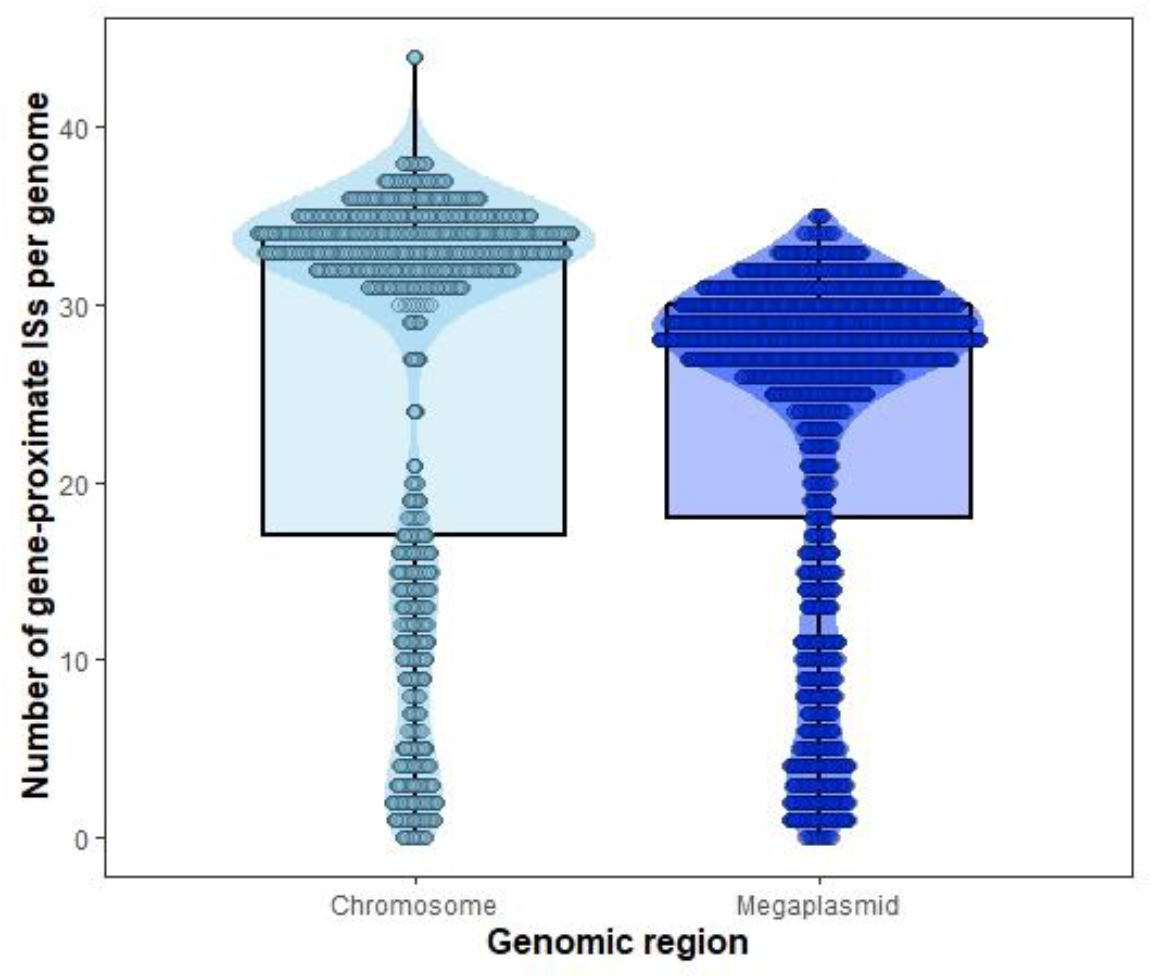
The megaplasmid contains significantly fewer gene-proximate IS than the chromosome. Boxplot showing number of IS that are close to (< 100 bp from start codon) or disrupt genes in the chromosome and the megaplasmid. Boxplots are coloured by genomic region. The difference in IS number between regions was tested statistically using a paired t-test.

**Figure S7.**
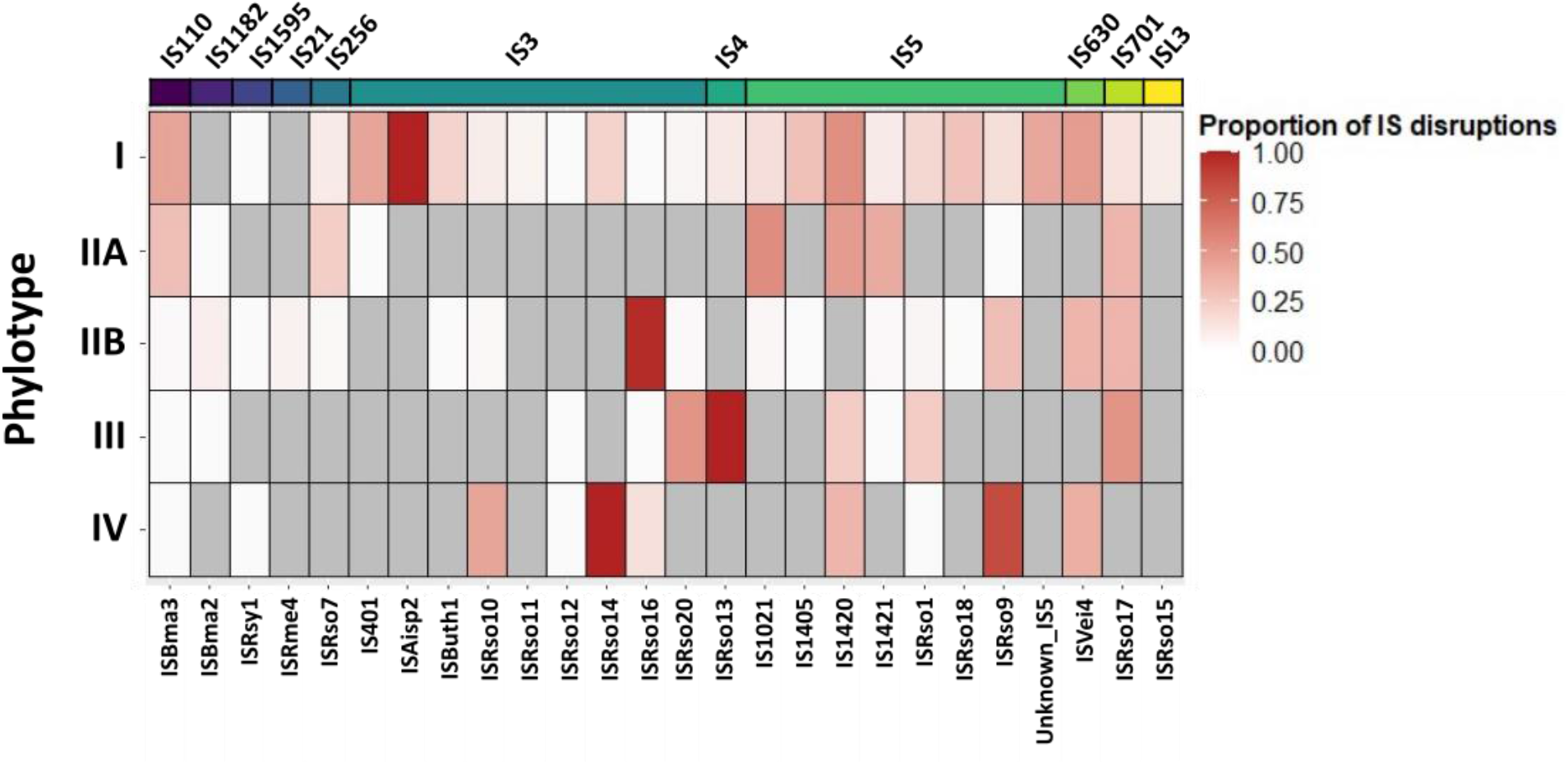
IS gene disruptions are caused by small number of IS subgroups and are lineage-specific. Heatmap showing the proportion of each IS subgroup that cause gene disruptions in each lineage. IS are clustered in the same way as Figure 2A, S4.

## Notes

### Competing Interest Statement

The authors have declared no competing interest.

